# Crystal Structure of Caryolan-1-ol Synthase, a Sesquiterpene Synthase Catalyzing an Initial Anti-Markovnikov Cyclization Reaction

**DOI:** 10.1101/2024.05.04.592530

**Authors:** Ramasamy P. Kumar, Jason O. Matos, Brandon Y. Black, William H. Ellenburg, Jiahua Chen, MacKenzie Patterson, Jacob A. Gehtman, Douglas L. Theobald, Isaac J. Krauss, Daniel D. Oprian

**Affiliations:** Department of Biochemistry, Brandeis University, Waltham, MA 02454; Department of Chemistry, Brandeis University, Waltham, MA 02454

## Abstract

In a continuing effort to understand reaction mechanisms of terpene synthases catalyzing initial anti-Markovnikov cyclization reactions, we solved the X-ray crystal structure of (+)-caryolan-1-ol synthase (CS) from *Streptomyces griseus*, with and without an inactive analog of the FPP substrate, 2-fluorofarnesyl diphosphate (2FFPP), bound in the active site of the enzyme. The CS-2FFPP complex was solved to 2.65 Å resolution and showed the ligand in a linear, elongated orientation, incapable of undergoing the initial cyclization event to form a bond between carbons C1 and C11. Intriguingly, the apo CS structure (2.2 Å) also had electron density in the active site, in this case density that was well fit with a curled-up tetraethylene glycol molecule presumably recruited from the crystallization medium. The density was also well fit by a molecule of farnesene suggesting that the structure may mimic an intermediate along the reaction coordinate. The curled-up conformation of tetraethylene glycol was accompanied by dramatic rotamer shifts among active-site residues. Most notably, W56 was observed to undergo a 90° rotation between the 2FFPP complex and apo-enzyme structures, suggesting that it contributes to steric interactions that help curl the tetraethylene glycol molecule in the active site, and by extension perhaps also a derivative of the FPP substrate in the normal course of the cyclization reaction. In support of this proposal, the CS W56L variant lost the ability to cyclize the FPP substrate and produced only the linear terpene products farnesol and α- and β-farnesene.

## INTRODUCTION

Terpenes form the largest class of natural products with over 80,000 unique members.^1–4^ They are characterized by ornate carbon skeleta with fused ring systems and multiple stereocenters, and are formed from achiral prenyl precursors in reactions which involve the generation and control of high-energy carbenium ion intermediates. The committed step in the biosynthesis of all terpenes is catalyzed by a terpene synthase.^5, 6^ Class 1 terpene synthases employ prenyl diphosphate substrates and use metal ions to trigger dissociation of the allylic diphosphate and formation of an initial resonance-stabilized carbocation.^5, 6^ Bond formation follows from the intramolecular reaction with electrons on double bonds present in the prenyl chain.^7, 8^ The reactions are terminated by either cation capture with a nucleophile such as water in the active site or by deprotonation to form an olefin.

Our interest in these reactions is currently focused on how some of the enzymes direct anti-Markovnikov attacks in which the carbocations form on secondary carbon atoms following cyclization instead of the more stable tertiary carbons.^9, 10^ Protein structures have been determined for several synthases that catalyze initial anti-Markovnikov cyclization reactions, including pentalenene synthase (PS)^10, 11^ and cucumene synthase,^9^ both from *Streptomyces*, trichobrasilenol synthase (TaTC6), a brasilane-type sesquiterpene synthase from the fungus *Trichoderma atroviride* FKI-3849,^12^ and the fungal enzymes presilphiperfolan-8β-ol synthase (BcBOT2) from *Botrytis cinerea*, Delta(6)-protoilludene synthase (DbPROS) from *Dendrothele bispora*, and longiborneol synthase (CLM1) from *Fusarium graminearum*.^13^ In the present work, we focus on enzymes from *Streptomyces.*^10^

Pentalenene synthase and cucumene synthase catalyze similar reactions, both resulting in the formation of triquinanes: an angular triquinane in the case of pentalenene synthase, and a linear triquinane in the case of cucumene synthase (Figure 1). Both reactions begin with displacement of the diphosphate moiety from carbon C1 of FPP, followed by addition of π-electrons from C11 to form the macrocyclic humulyl cation A, with the carbocation located on C10. Subsequent hydride shift from C9 to C10 results in migration of the positive charge to C9, which is required for transannular cyclization in the next step to form the first fused 5-carbon ring of the triquinane scaffold. An X-ray crystal structure for PS in complex with an inactive difluoro-farnesyl diphosphate substrate analog (DFFPP) shows the ligand to be coiled in the active site of the enzyme, with the reactive C1 and C11 carbons separated by only 4.0 Å, poised in a ready-to-react conformation.^10^ Critically, the C9 carbon is seen to be located 3.5 Å directly above the center of the phenyl ring in the side chain of F76, in perfect position to facilitate the hydride shift through C-H…π interaction and stabilize the resulting positive charge on C9 through cation-π interaction.^14–21^ In this way, the enzyme directs regioselectivity in the initial anti-Markovnikov cyclization reaction, stabilizing the positive charge on C9 of humulyl cation B. This arrangement of atoms in the active-site complex suggests the possibility that the initial cyclization reaction is concerted, with diphosphate release, C1/C11 bond formation, and C9 to C10 hydride shift all taking place simultaneously. While a structure for cucumene synthase with ligand bound has not yet been reported, it seems likely that it would look similar to that of PS-DFFPP, and it is noteworthy that cucumene synthase also has an aromatic residue (Tyr) at the same position as F76 in PS.^10^

**Figure 1.**
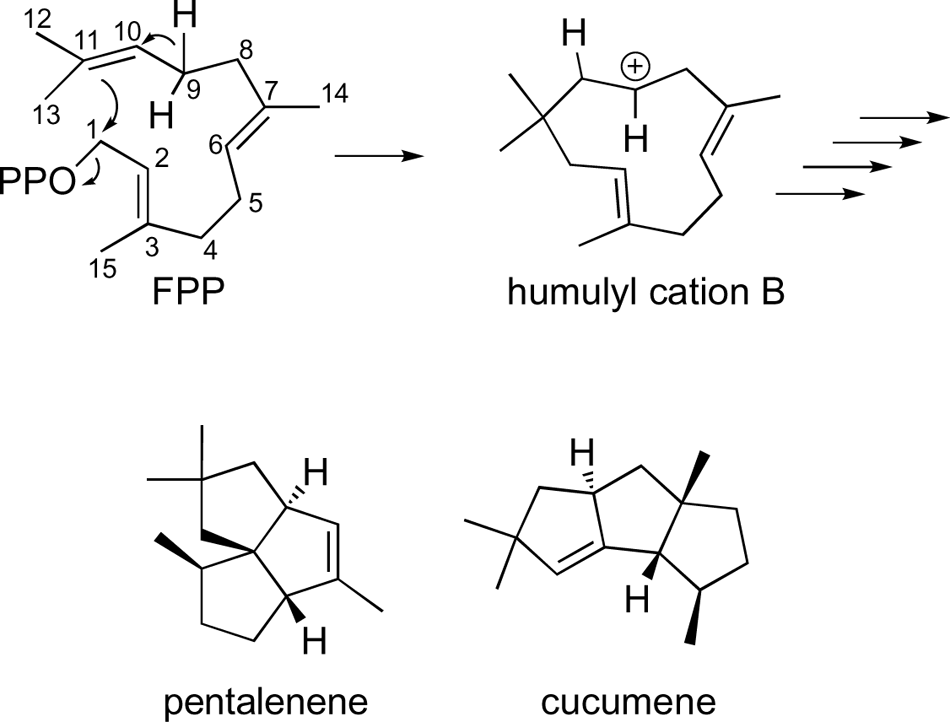
Initial cyclization reactions for the enzymes PS and cucumene synthase showing intermediate formation of humulyl cation B.

We report here structures for the apo and ligand-bound forms of a third terpene synthase from *Streptomyces* that catalyzes an initial anti-Markovnikov cyclization reaction, the enzyme caryolan-1-ol synthase (CS).^22^ CS is a 38 kDa monomer from *Streptomyces griseus* that is composed of a single domain with typical class I α-helical terpenoid cyclase fold and contains conserved Asp-rich (DDXXD) and NSE/DTE motifs for binding of the diphosphate moiety of the substrate along with three Mg^2+^ ions.^5^ The CS reaction (Figure 2) is of interest because it begins with release of the diphosphate and attack of the C11 π-electrons on C1 to form humulyl cation A, with carbocation located on C10, as is the case with both PS and cucumene synthase, but the reaction does not continue with a hydride shift and migration of the positive charge to C9. Instead, π-electrons on C2 undergo transannular attack on C10 with formation of a fused 4-carbon ring. Intriguingly, CS does not have an aromatic residue at the position corresponding to F76 in PS. Instead, CS has an Ala at this position (A79), accounting for the absence of a hydride shift in the reaction. Surprisingly, and despite obvious similarities in the reactions catalyzed by the two enzymes, we find significant differences in the biochemical behavior and X-ray crystal structure of CS in comparison with PS such that the effect of active site mutation and conformation of bound ligands in CS are not easily predicted from results with PS. As a consequence, these results suggest that there are fundamental differences in the reaction mechanisms for these two enzymes.

**Figure 2.**
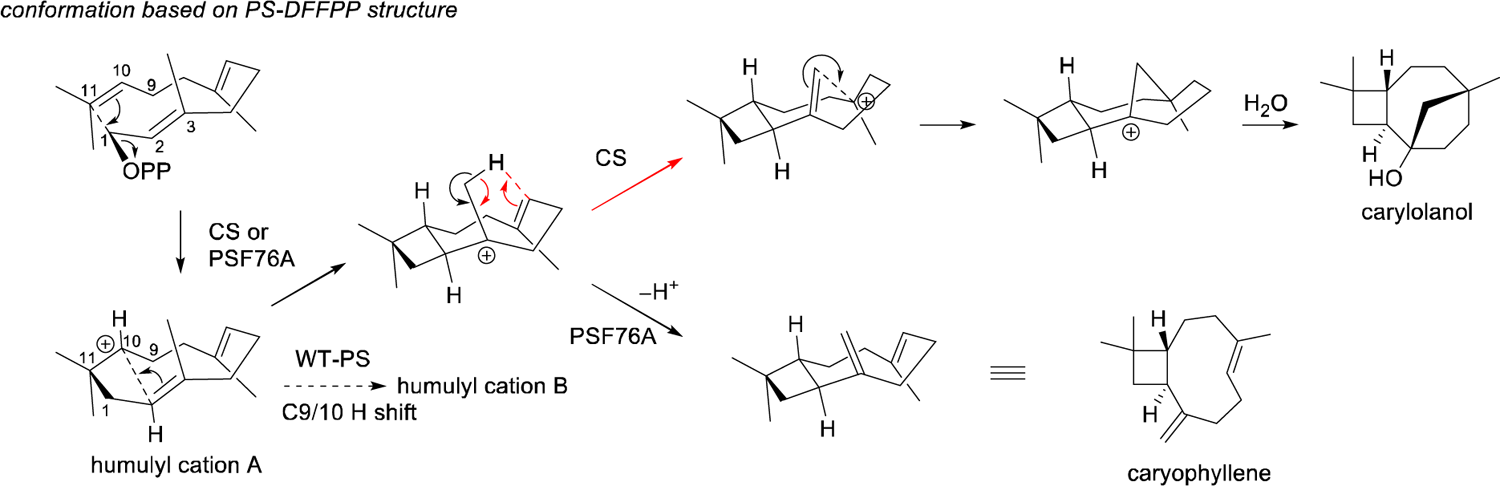
Reaction scheme for CS. The reaction begins with the FPP substrate, shown in the upper left corner of the figure with carbon atoms numbered. Initial anti-Markovnikov cyclization involves loss of the allylic pyrophosphate from C1 to form a resonance-stabilized farnesyl cation and attack of π electrons from C11 on C1 with resulting carbocation on C10. Subsequent transannular cyclization steps and addition of water result in the multicyclic fused ring system of caryolan-1-ol.

## RESULTS

### F76A Variant of Pentalenene Synthase

The F76A variant of PS was constructed to determine how the reaction would be modified by removal of the aromatic side chain with substitution of an Ala for Phe at this position. As is shown in Figure 3A, overnight incubation of the variant with FPP produced a significantly modified product profile with near complete loss of the normal product pentalenene and the appearance of several new species in a promiscuous product profile dominated by caryophyllene. Caryophyllene arises as a consequence of carbocation development on C10, as is the case with the CS reaction (Figure 2), and not on C9 as is the case with PS WT. Initial rate assays in Figure 4A show that the mutation causes an estimated 38-fold decrease in *k*_cat_ as monitored by the change in summed areas under the peaks of the GC profiles, while the production of pentalenene is essentially eliminated. These data support the conclusions of Matos et al.^10^ that F76 directs regioselectivity for development of a carbocation on C9 in the initial anti-Markovnikov cyclization reaction catalyzed by PS WT.

**Figure 3.**
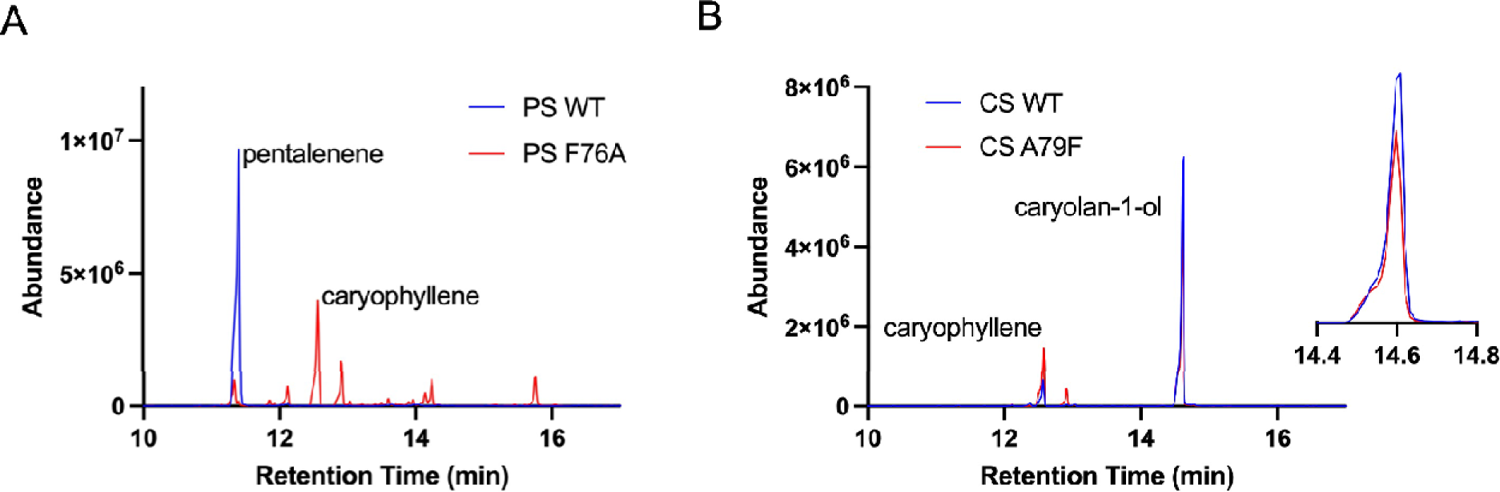
GC-MS product profile for overnight reaction of FPP with WT and variants of PS and CS. (A) Reaction with PS WT and variant PS F76A. (B) Reaction with CS WT and variant CS A79F. Inset shows the peaks for caryolan-1-ol with expanded time axis to allow for better resolution of the red and blue peaks.

**Figure 4.**
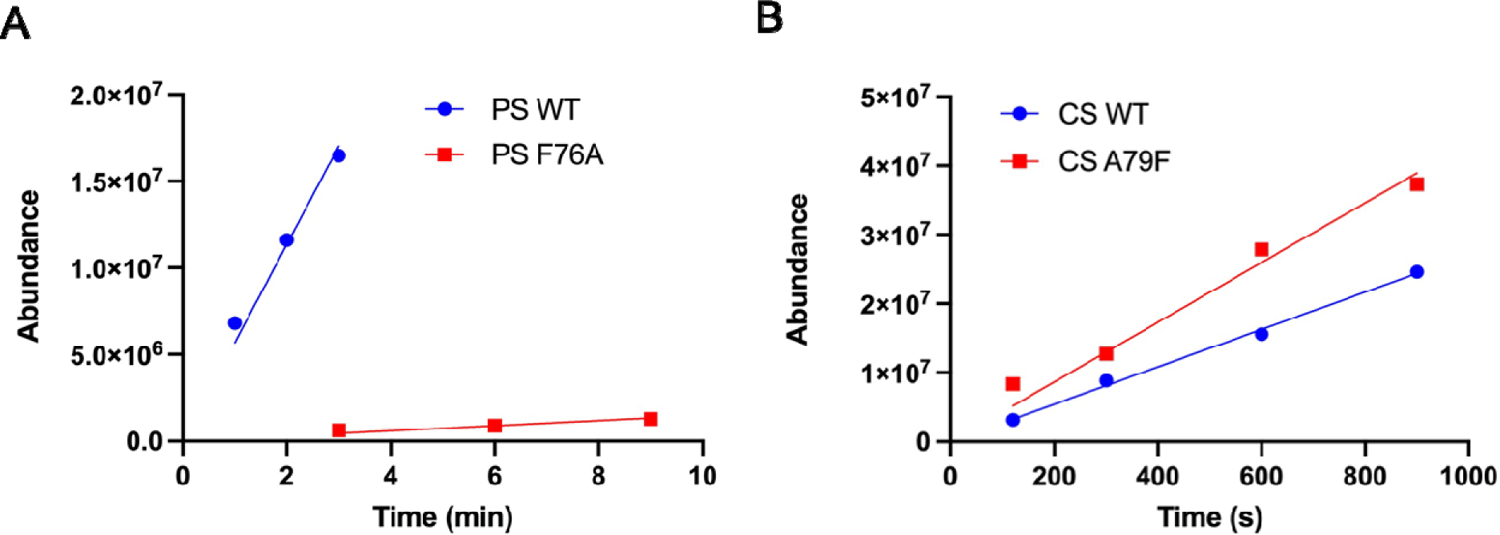
Initial rate data for reactions of PS and CS. The reactions were performed under V_max_ conditions for: (A) PS WT and the PS F76A variant; and (B) CS WT and CS A79F variant.

When the complementary mutation is made in CS (A79F) the result is very different from that observed for PS. Notably, the CS reaction is little affected by the mutation: the product profile is hardly changed from that of the CS WT reaction (Figure 3B), and the reaction rate is not observed to be significantly altered from that of the WT reaction (Figure 4B).

### Structure of CS

To further explore the underlying differences between the PS and CS enzymes, crystals of apo CS were grown in the presence of CaCl_2_ and PEG400 precipitants. The structure was determined by X-ray diffraction in space group *P*2_1_ to 2.3 Å resolution using the MoRDa automatic molecular replacement pipeline.^23^ There are four molecules in the asymmetric unit, each of which displays the typical 11-helix “terpene synthase fold” as observed for PS and other sesquiterpene synthase enzymes.^5, 10, 11^ Similar patterns of missing electron density were observed for the protein in all four molecules for the N-terminus up to E9, part of the DDxxD motif, F225-E234 of the polypeptide, and the C-terminus beyond R311.

As shown in Figure 5A, a crescent-shaped electron density was observed in the active site of all four molecules of the asymmetric unit. This density is surrounded by amino acids W56, A79, F80, D83, W148 (not shown), and F183 in the active site core and was well fit by a single molecule of tetraethyleneglycol, presumably recruited from the crystallization medium.

**Figure 5.**
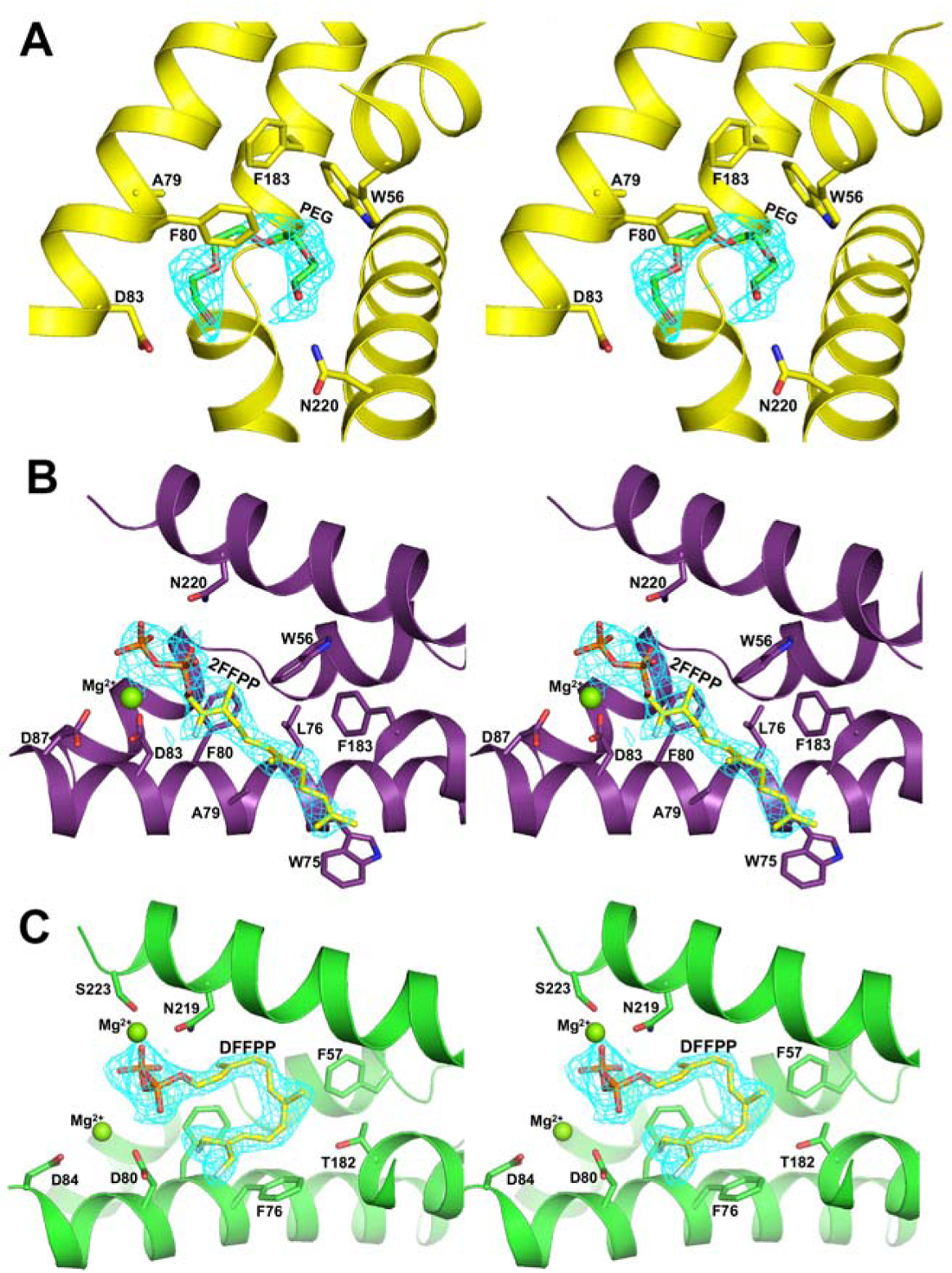
Electron density (*F_o_ – F_c_* = 3σ cutoff colored in cyan) shown in stereoview for ligands in the active sites of CS and PS. (A) Electron density found in the active site of apo CS (yellow). A tetraethyleneglycol molecule (H(C_2_H_4_O)_4_OH; molecular mass = 194) is shown modeled into the electron density. (B) Electron density found in the active site of CS (purple) that had been co-crystallized with 2FFPP. 2FFPP is shown modeled into the electron density. (C) Electron density found in the active site of PS (green) after crystals had been soaked with DFFPP. DFFPP is shown modeled into the electron density. Mg^2+^ ions in panels (B) and (C) are shown as green spheres. The orientation of CS and PS is similar in panels (B) and (C).

### Structure of CS bound to the substrate analog 2FFPP

The 2FFPP substrate analog was shown to be an effective inhibitor of the CS reaction (Figure 6) and was selected for further studies aimed at determination of the ligand-bound complex. Crystals of CS bound to 2FFPP were obtained by co-crystallization under the same conditions as had been optimized for the apo-enzyme. The space group and cell dimensions were the same as observed for the apo-enzyme, and the structure was determined by molecular replacement to 2.65 Å resolution using the apo-enzyme as a search model. After initial refinement, a difference Fourier map showed electron density corresponding to the ligand extending well into the active site in each of the molecules of the asymmetric unit. The 2-fluoro substrate analog 2FFPP was modeled into the four molecules independently. While the prenyl chain adopted a similar conformation in each of the active sites, minor differences were noted. For example, a Mg^2+^ ion was observed only in molecules A, C, and D, and the diphosphate moiety exhibited a different orientation in each of the active sites. Molecule A displayed the best resolved electron density for the ligand, Mg^2+^ ion, and DDxxD motif compared to the other molecules and, therefore, was used for the following analysis (Figure 5B).

**Figure 6.**
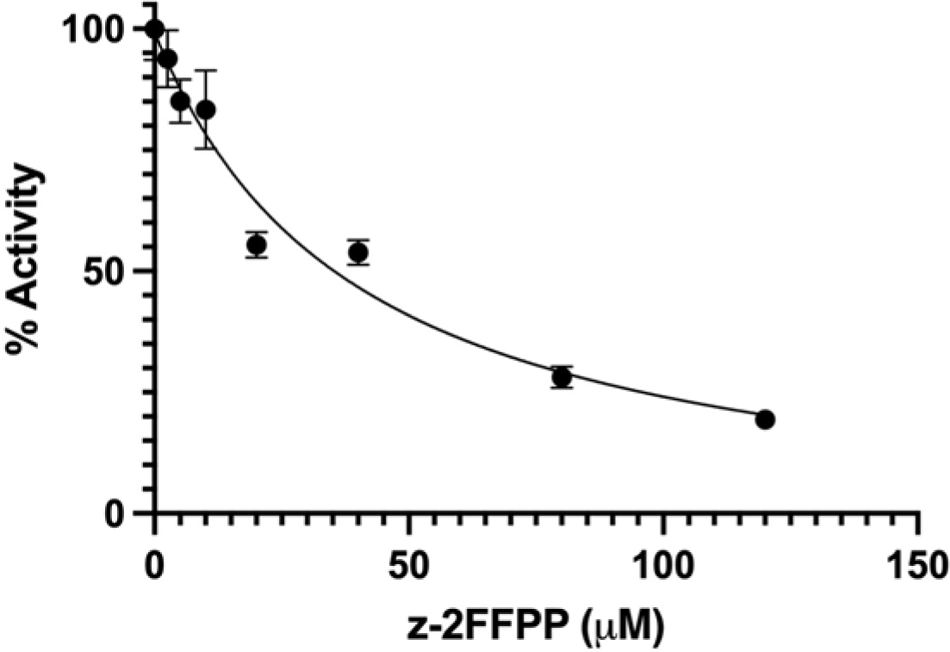
Inhibition of the CS reaction by 2FFPP. Each data point was determined from initial rates measured with 1μ CS and 50 μM FPP. Error bars are standard deviations.

The 2FFPP ligand is in a linear, extended conformation (Figure 5B) and not in the reaction ready conformation observed previously for the PS-DFFPP complex (Figure 5C), where the prenyl chain curves in the active site to position carbon atoms C1 and C11 close to each other in position to undergo the initial anti-Markovnikov cyclization reaction.^10^ Interestingly, several amino acid residues in the active site of CS appear to undergo movement upon binding of the 2FFPP ligand (Figure 7), reminiscent of the induced-fit mechanism described for selinadiene synthase.^24^ A179 undergoes a large 2.6 Å movement to accommodate the C15 methyl group of the 2FFPP ligand, causing backbone rearrangement of residues 177-184 in the G1-G2 helix region. Steric hindrance from the prenyl chain causes the side chain of F183 to rotate by approximately 90° and move 2.7 Å towards W56 which in turn causes a 90° angle rotation of the W56 side chain. Interestingly, W56 and F183 appear to be the residues primarily responsible for the curved conformation observed for the tetraethyleneglycol molecule found in the active site of the apo CS structure. Finally, residues 87-93 are partially disordered in apo CS but become fully ordered upon binding of the ligand and metal ion, bringing D87 of the DDxxD motif into position to coordinate with the active site Mg^2+^.

**Figure 7.**
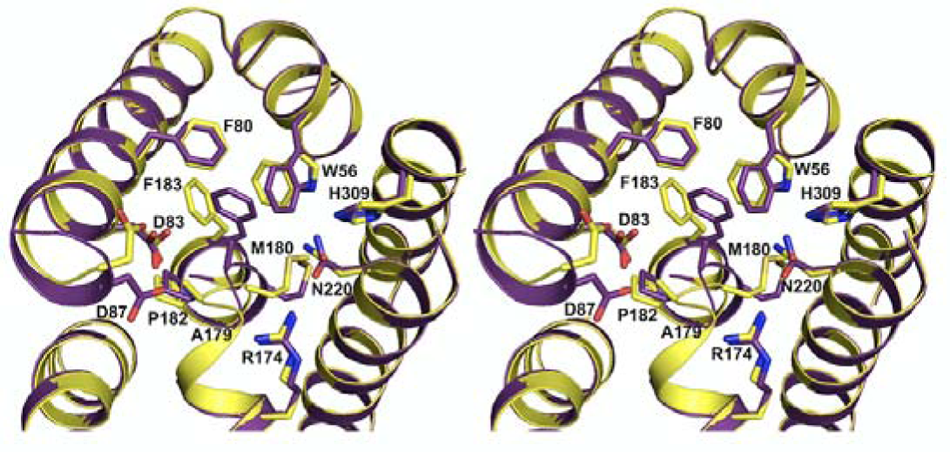
Superposition of active sites of apo (yellow) and 2FFPP-bound CS (purple) in stereoview showing the conformational change of amino acid side chains triggered by the binding of 2FFPP. The ligands PEG and 2FFPP were omitted for clarity.

### CS W56L

The large rotamer shift observed for W56 associated with binding of the 2FFPP and the proximity to the curved density for PEG in the active site made this residue of interest to target for mutagenesis. We chose to replace the Trp residue with Leu as Leu is often observed at this conserved position in sesquiterpene synthases from *Streptomyces*. As is shown in Figure 8, the CS W56L variant displayed a dramatically altered product profile which was dominated by the linear products farnesol and α- and β-farnesene, suggesting that the enzyme has lost the ability to cyclize the substrate molecule.

**Figure 8.**
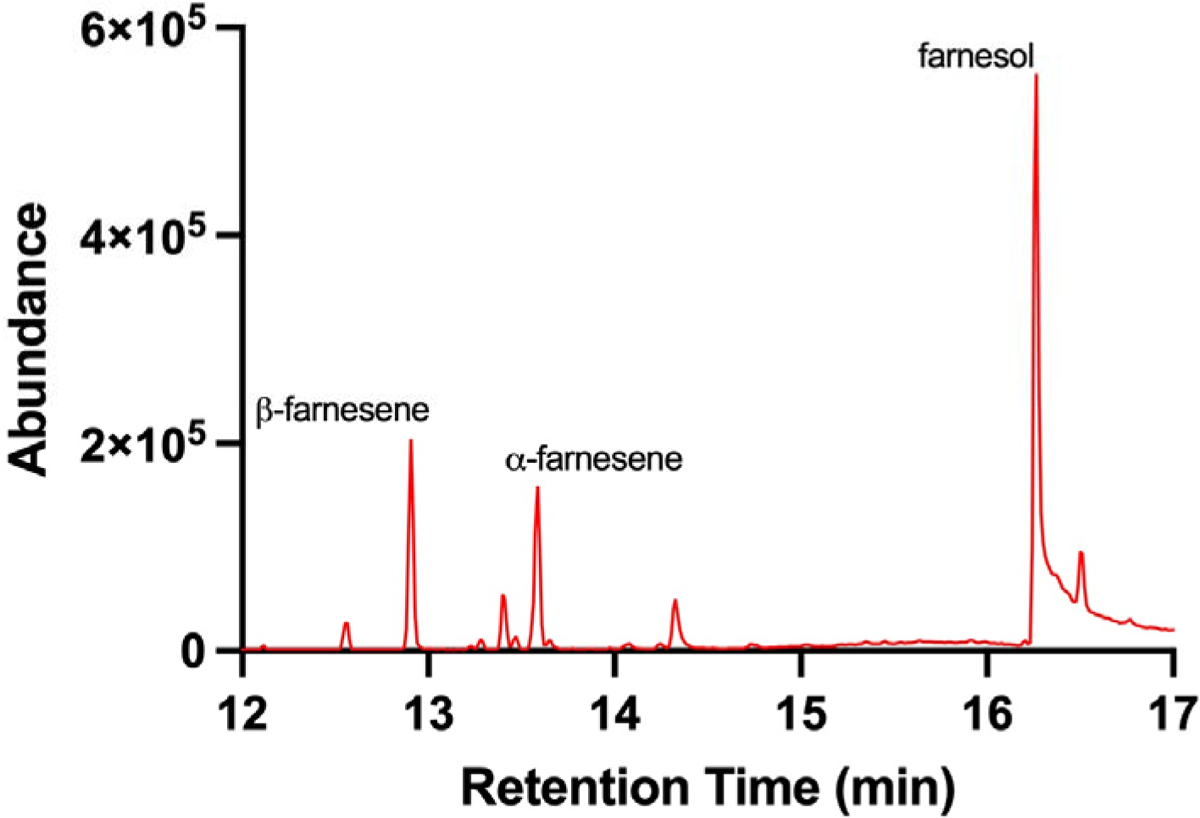
GC-MS product profile for overnight reaction of FPP with CS W56L. The product profile for the variant is predominantly composed of the linear products farnesol and α- and β-farnesene.

### Structure of PS F76A

Apo PS F76A was crystallized from PEG400 and ammonium sulfate solutions in space group *P*6_3_ with two molecules in the asymmetric unit. The structure was determined to 2.5 Å resolution by molecular replacement using the PS-DFFPP complex as a search model (PDB entry 6WKD). The first five residues of the N-terminal segment, and residues beyond S311 (in molecule A; beyond D316 in molecule B) in the C-terminus were not resolved in the electron density. The overall structure of the protein is very similar to the PS-DFFPP complex reported earlier.^10^ The active site pockets for both molecules in the asymmetric unit have an electron density similar to that described above for apo CS (compare Figures 5A and 7A), and the density is well fit in apo PS with a tetraethyleneglycol molecule as was the case also for apo CS. The conformation of the tetraethyleneglycol molecule appears to be guided by the side chains of amino acids F57, M73, A76, F77, D80, T182, and W308 in the active site pocket (T182 and W308 were omitted from the figure for clarity).

### Structure of PS F76A ligand complexes

Crystals of apo PS F76A grown under similar conditions to those previously reported for the PS WT^10^ were soaked with 1 mM DFFPP or 2FFPP in the presence of 10 mM MgCl_2_ and 15% glycerol cryoprotectant. The structures of DFFPP- and 2FFPP-bound PS F76A were determined to 2.65 Å and 2.2 Å resolution, respectively, using WT-DFFPP (PDB entry 6WKD but without the ligand) as a search model for molecular replacement. In both enzyme complexes, electron density for a few residues in the N- and C-terminal regions were not resolved and were not included in the final model. Clear electron density was observed for the ligands in molecule A of the asymmetric unit along with three Mg^2+^ ions in complex with the diphosphate moiety. Structures for both DFFPP and 2FFPP (Figure 9B and C) were clearly observed to be oriented in a clockwise direction in the active site from the perspective of the opening to the active site at the surface of the protein. This is in contrast to the previously reported structure for PS WT where the ligand is curled in a counterclockwise direction and C1 and C11 are positioned within 4.0 Å ready to undergo the initial cyclization reaction. C1 and C11 are further apart in both of the F76A structures (6.1 Å for 2FFPP; 5.3 Å for DFFPP), and the C14 methyl group of the respective ligand occupies the void left by the phenyl side chain of F76.

**Figure 9.**
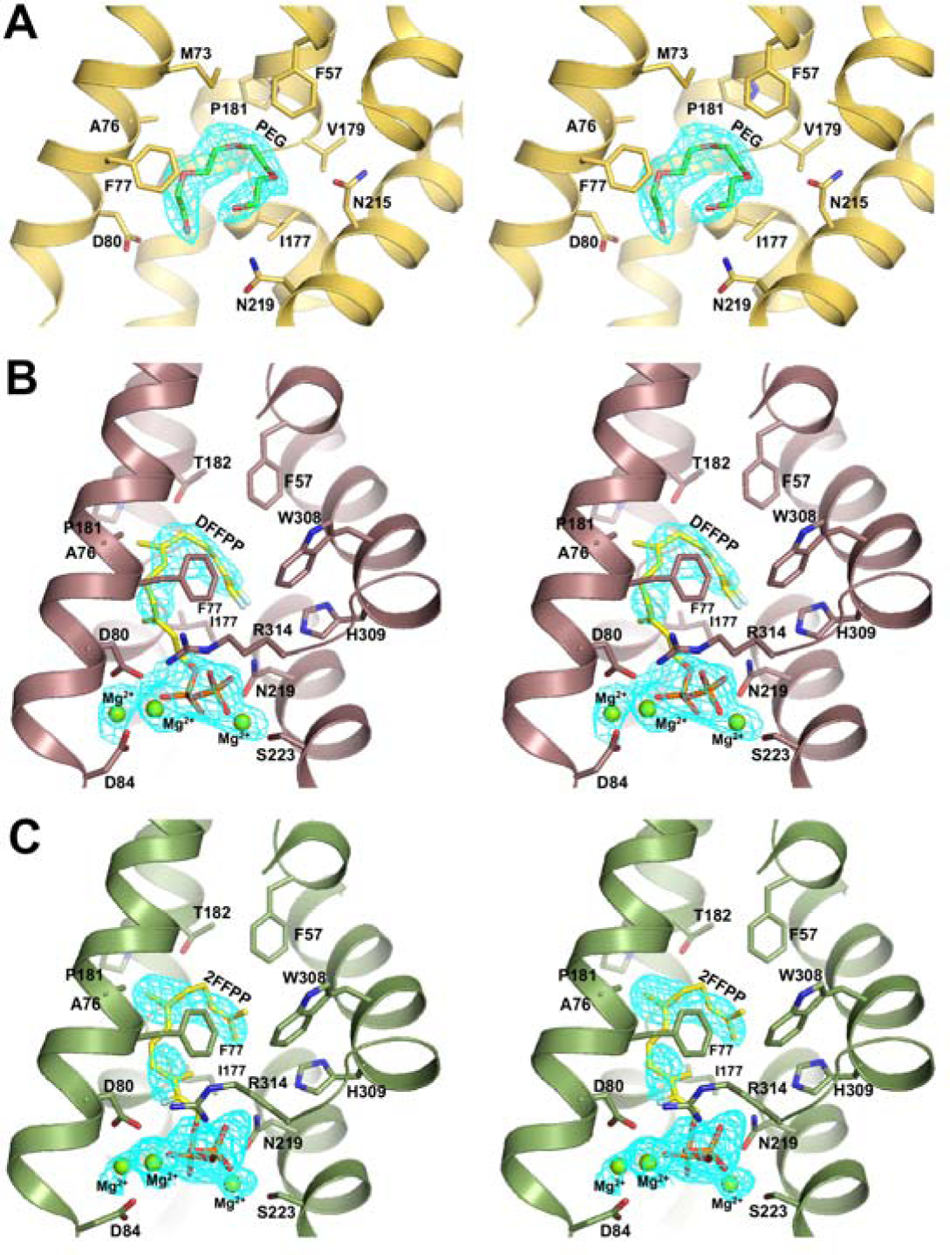
Electron density (*F_o_ – F_c_* = 3σ cutoff colored in cyan) shown in stereoview for ligands and Mg^2+^ ions in the active site of PS F76A. (A) Electron density found in the active site of apo PS F76A (yellow). A PEG molecule (H(C_2_H_4_O)_4_OH; molecular mass = 194) is shown modeled into the electron density (green). (B) Electron density found in the active site of PS F76A (brown) after crystals of the apoprotein had been soaked with DFFPP. DFFPP (yellow) is shown modeled into the electron density. (C) Electron density found in the active site of PS F76A (green) after crystals had been soaked with 2FFPP. 2FFPP (yellow) is shown modeled into the electron density. Mg^2+^ ions in panels (B) and (C) are shown as green spheres. The orientation of PS F76A is similar in panels (B) and (C).

## DISCUSSION

This study was motivated by our previous work on the anti-Markovnikov cyclization reaction catalyzed by PS in which regioselectivity in directing carbocation development on the secondary carbon C9 of the FPP substrate is governed by the aromatic side chain of F76.^10^ We were interested in the CS reaction because the initial cyclization reaction also involves attack of C11 on C1 to displace the diphosphate moiety and produce a carbocation on C10 of the substrate, but the CS reaction does not involve a subsequent hydride shift with development of positive charge on C9. Intriguingly, CS has an alanine (A79) at the corresponding position to F76 in PS, removing the possibility of π-cation stabilization of positive charge on C9 as a result of interaction with an aromatic side chain. And indeed, the F76A variant of PS displays a dramatic loss of activity (Figure 4A) and associated change in product profile which is dominated by caryophyllene (Figure 3A), the product expected from a carbenium ion on C10, and not C9. Somewhat disappointingly, the complementary mutation in CS A79F was almost without effect; there was very little change in activity (Figure 4B) or product profile (Figure 3B), suggesting a fundamental difference in the enzymatic reactions catalyzed by PS and CS.

This conclusion was supported by the crystal structure for CS bound to the inactive substrate analog 2FFPP in which the ligand is bound to the enzyme in an extended conformation with carbon atoms C1 and C11 separated by 11.3 Å (Figure 5B), not in a ready-to-react conformation, as is the case for DFFPP in PS (Figure 5C), indicating that the CS reaction starts with a non-productively bound ligand that must undergo a conformational change to position C1 and C11 close enough to form a covalent bond in initial cyclization of the FPP substrate. In this context, it is interesting that the apo CS structure has electron density in the active site corresponding to a crescent-shaped conformation of tetraethyleneglycol which was likely recruited from the crystallization medium. The PEG400 precipitant used in the crystallization conditions has a range of sizes extending from 200 to 600 g/mol with average at 400 g/mol,^25^ emphasizing that the enzyme must have specifically recruited one of the least abundant molecules (194 g/mole) into the active site. This may not be so surprising considering that the PEG electron density is also very well fit by farnesene, which has the carbon skeleton of the FPP substrate following dissociation of the diphosphate moiety (Figure 10). The distance between C1 and C11 in this model (6.1 Å) is dramatically decreased from that for 2FFPP in its extended conformation. It is tempting to speculate that the density observed in the active site of apo CS may mimic an intermediate in the reaction coordinate that follows initial binding of an extended conformation of the prenyl chain, perhaps requiring dissociation of diphosphate and formation of a farnesyl cation (Figure 2), in a more stepwise mechanism, to trigger the change. This hypothesis is further bolstered by the fact that mutating W56 to a Leu completely abolishes CS’s ability to form cyclic products. W56 is one of the residues in the active site which undergoes a large rotamer shift between the apo and ligand-bound structures and appears to be a residue largely responsible for the curled conformation of the tetraethylene glycol molecule. Without the aromaticity or steric bulk of Trp, it appears that CS can no longer achieve this substrate orientation. PS F76A also binds both 2FFPP and DFFPP in non-productive conformations (Figure 9B and C), which may account in part for the loss of activity observed for this variant.

**Figure 10.**
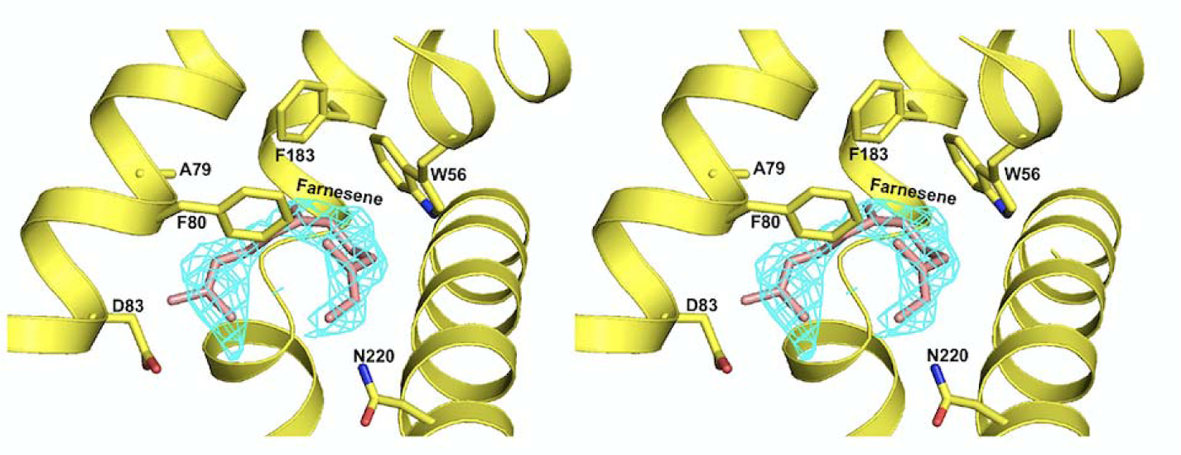
Active-site of CS (yellow) in stereoview showing electron density for tetraethyleneglycol (*F_o_ – F_c_* = 3σ cutoff colored in cyan) modeled with a molecule of farnesene (tan).

Despite similarities in the underlying chemistry, PS and CS clearly have significant differences in their respective enzymatic reaction mechanisms, and one wonders whether these differences appear more generally in the large family of terpene synthases. PS and CS locate to different clades in the TS phylogenetic tree,^26^ and it is of interest to determine if the difference in mechanism tracks within each clade or not. Future studies will focus on this question.

## METHODS

### Preparation of tris-ammonium (2Z,6E)-2-fluoro farnesyl pyrophosphate

The synthesis of 2FFPP was performed as described by Miller et al. (2003)^27^ except that phosphorylation of the 2-fluoro farnesol intermediate was performed by the method of Keller and Thompson (1993)^28^, as we have previously described.^10, 29^

### Preparation of CS, CS A79F, and PS F76A

The genes for CS from *Streptomyces griseus* (B1W019.1)^30^ and CS A79F were codon optimized for expression in *Escherichia coli* and included a glycine codon immediately following the start methionine. The CS WT and CS A79F genes were cloned into a pET-28a vector as a NcoI and EcoRI cassette for expression in *E. coli*. A His_6_ tag and TEV protease cleavage site were also included at the N-terminus of the protein.

The plasmids were transformed into BL21 (DE3) competent cells, and colonies were selected after growth on 50 μg/mL kanamycin containing LB/agar plates. A single colony was used to inoculate 10 mL of LB broth containing 50 μg/mL kanamycin. This suspension was incubated with shaking (220 rpm) at 37°C overnight. The entire culture was then used to inoculate 1 L of LB broth containing 50 μg/mL kanamycin. Once an optical density of 0.6-0.8 was reached, expression was induced by addition of 500 μM IPTG, and incubation continued with shaking at 16°C overnight.

CS WT and CS A79F were purified as follows. Cells were harvested by centrifugation and resuspended in 50 mM Tris buffer, pH 8, containing 150 mM NaCl and 10% glycerol (Buffer A). The suspension was sonicated (30-40 Watts, 4 min total, 20 s on, 40 s off), and the supernatant fraction clarified by centrifugation. The resulting lysate was filtered through a 0.2 μm nylon filter and loaded onto a prepacked 5 mL HiTrapFF Ni-Sepharose column (GE Healthcare Life Sciences) that had been equilibrated in Buffer A. The column was then washed with 5 column volumes of Buffer A containing 5 mM imidazole, and protein was eluted with a linear gradient of 5 mM to 1 M imidazole in Buffer A. Fractions were collected in 50 mL Falcon tubes with 10 mL of Buffer A to dilute the imidazole. Imidazole was then removed from the solution and the protein concentrated by repeated cycles of concentration and dilution in Buffer A using Amicon 30 kDa cutoff Ultra −15 filtration units. Concentrated protein was aliquoted and flash frozen using liquid nitrogen before storing at −80°C until further use.

Due to the inherent instability of the PS F76A variant, purification of the protein was essentially the same as for CS WT and CS A79F described above except that the procedure began with 4 L of culture, and Buffer A was replaced with a solution of 50 mM Bis-Tris propane, pH 7.5, containing 150 mM KCl, 10 mM MgCl_2_, and 10% glycerol (Buffer B).

### Enzymatic Assays

Enzymatic assays were performed as previously described in Matos et al. (2020).^10, 31^ Hexane extractable products were identified and quantified using GC-MS (Agilent Technologies 7890A GC system coupled with a 5975C VL MSD with a triple axis detector). Pulsed-splitless injection was used to apply 5 μL samples to a HP-5 ms (5%-phenyl)-methylpolysiloxane capillary GC column (Agilent Technologies; 30 m X 250 μm X 0.25 μm) at an inlet temperature of 220°C and a transfer temperature of 240°C, run at constant pressure using helium as a carrier gas. Samples were initially held at 50°C for 1 minute, followed by a linear gradient of 10°C/min to 220°C, which was then held for 10 minutes.

Two categories of assays were performed: the first was overnight assays to determine the total product profile of each enzyme; the second was initial rate assays performed to calculate activity of the enzymes. The overnight assays were performed in Buffer A with 10 mM MgCl_2_ for CS WT and CS A79F while PS F76A assays were performed in Buffer B. Enzyme concentration was 1 μM, and the reaction initiated by addition of 200 μM FPP. Immediately after initiation of the reaction, 1 mL of hexanes was layered on top of the reaction solution.

Following incubation at room temperature overnight, the reactions were quenched by vigorous vortexing, extracting products into the hexane layer which was then analyzed by GC-MS. Initial rates were determined in a similar manner, except that the enzyme concentrations varied depending on which enzyme was being used, and early time points were analyzed to determine initial rates for the reactions. PS WT and PSF76A assays were performed at an enzyme concentration of 100 nM, while CS WT and CS A79F assays were performed at an enzyme concentration of 1 μM. PS assays were initiated with 50 μM FPP while CS assays were initiated with 100 μM FPP.

### Inhibition Assays

Inhibition of CS with 2FFPP was performed by the same procedure described above for initial rates except that the reactions contained 50 μM FPP and a variable amount of 2FFPP (0-200 μM). In addition, reactions were initiated by addition of enzyme.

### Crystallization, Data Collection, Processing, and Refinement

#### CS

7.5 mg/ml of protein in 50 mM Tris, 150 mM NaCl and 10% glycerol was mixed with an equal volume of precipitant from a Hampton sparse matrix screen (Hampton Research, Aliso Viejo, CA) in sitting drop vapor diffusion crystallization trays using a Gryphon robot (Art Robbins Instruments, Sunnyvale, CA). Crystals, which appeared in about two weeks, were frozen in the presence of 15% glycerol. Data sets were collected at beamline 8.2.1 at the Advanced Light Source (Lawrence Berkeley National Laboratory, Berkeley, CA). A crystal grown in 28% PEG400, 200 mM CaCl_2_ and 100 mM HEPES, pH 7.5, diffracted to 2.3 Å resolution and was used for analysis of the apo-enzyme structure. The data set was auto-processed in XIA2^32^ and scaled using SCALA^33^ from the CCP4 suite^34^. Diffraction data were processed in the *P*2_1_ space group with cell dimensions *a* = 68.5 Å, *b* = 89.8 Å and *c* = 114.1 Å, and *α* = *β* = 90° and *γ* = 90.8°. The CS structure was solved from CS sequence and structure factors introduced into the MoRDa automatic molecular replacement pipeline^23^ available from CCP4 online^35^. The pipeline server picked apo-hedycaryol synthase (PDB 4MC0) as a search model. The output model was automatically uploaded to the ARP/wARP online server^36, 37^ where the model was built to a score of 0.51. The score was improved to 0.83 with another 50 rounds of rebuilding resulting in a R and R_free_ value of 0.211 and 0.454, respectively, for 10277 atoms including dummy atoms. The resulting model was improved further using phenix.autobuild^38^ from the PHENIX suite^39^ and was refined with phenix.refine^40^ to R and R_free_ values of 0.216 and 0.275, respectively. Missing amino acids resolved up to *2F_o_ – F_c_* = 1σ in all four molecules of the asymmetric unit were built using coot^41^ followed by another round of phenix.refine. At this stage we clearly saw difference Fourier electron density (*F_o_ – F_c_* = 3σ) in the active site of each molecule in the asymmetric unit and modelled it with a molecule of tetraethyleneglycol. Six Ca^2+^ ions with coordinated water molecules were also modeled. The final round of refinement included all water molecules and had R and R_free_ of 0.21 and 0.256, respectively.

#### CS-2FFPP

The crystals of CS in complex with the 2FFPP substrate analog were obtained by co-crystallization using the same crystallization conditions as was used for the apo-enzyme but including 1 mM 2FFPP and 10 mM MgCl_2_. Several crystals appeared and the best diffracted to 2.65 Å resolution. The same beamline, space group, and similar cell dimensions as apo CS were used to autoprocess the data with XDS^42^. Data were scaled using SCALA^33^ from the CCP4 suite^34^. The structure was solved by molecular replacement using PHASER^43^ from CCP4^34^ with apo-CS as a search model. After initial rounds of refinement, the active site of all four molecules in the asymmetric unit contained an extended, linear electron density accompanied with characteristic metal ion peaks. Slight differences were observed in each active site, and the ligands were modelled independently as guided by electron density in each of the molecules. We also found a few free phosphate ions at protein interfaces between molecules in the asymmetric unit. The final model was refined to R and R_free_ of 0.204 and 0.249, respectively.

#### PS F76A

The crystal used in solving the structure of the PS F76A apo-enzyme was obtained from 100 mM HEPES, pH 7.5, containing 2 M ammonium sulfate and 2% PEG400 precipitants. Crystals of PS F76A in complex with the 2FFPP and DFFPP ligands were prepared by soaking apo PS F76A crystals with the substrate analog, but in this case the apo PS F76A was crystallized using the same conditions reported previously for the WT enzyme (0.8−1.2 M sodium tartrate, 100 mM Tris, pH 7.5− 9, and 5 mM DTT). All crystals were frozen in the presence of reservoir solution containing 15% glycerol.

Diffraction data sets were collected at the Northeastern Collaborative Access Team (NE-CAT) beamline 24-ID-E at the Advanced Photon Source (Argonne National Laboratory, Lemont, IL) using a Dectris EIGER 16M detector (DECTRIS Ltd.). The best crystals of PS F76A, PS F76A-2FFPP, and PS F76A-DFFPP diffracted to 2.5 Å, 2.2 Å, and 2.65 Å, respectively. The data were autoprocessed using XDS^42^ and scaled using SCALA^33^ from the CCP4 suite^34^. The structures were solved in space group *P*6_3_ by molecular replacement using PHASER^43^ with molecule A of the PS-DFFPP complex (PDB entry 6WKD) without ligand and solvent molecules as a search model. The solution for all data sets had two molecules in the asymmetric unit. After an initial 10 cycles of refinement, there was clear difference Fourier electron densities above 3σ cut off in a crescent shape seen in the active sites of both molecules A and B of apo PS F76A, similar to that described for apo CS above, and the density was well fit by a molecule of tetraethyleneglycol. Electron densities for the substrate analogs and associated metal ions in the soaked crystals were observed only in the active sites of molecules A. Final models were refined to R/R_free_ of 0.211/0.239, 0.192/0.215, and 0.198/0.229 for apo PS F76A, PS F76A-2FFPP, and PS F76A-DFFPP, respectively.

All five data sets were refined with rigid-body refinement followed by positional and B-factor refinement using phenix.refine^40^ from the PHENIX software suite version 1.20^39^. Simulated annealing was included in earlier refinements to minimize the initial model bias. Manual model building was performed using COOT version 0.9. All ligands were modeled using Jligand version 1.0 from CCP4 software suite version 8.0^34^, and the generated coordinates and restraints were used for further refinements. Solvent molecules were searched for and included in the final round of refinement. The amino acids and solvent molecules not resolved above *2F_o_ – F_c_*= 1σ were not included in the final models. Complete data collection statistics are listed in Table 1. Data sets for CS (PDB entry xxxx), CS-2FFPP (PDB entry xxxx), PS F76A (PDB entry xxxx), PS F76A-2FFPP (PDB entry xxxx), and PS F76A-DFFPP (PDB entry xxxx) have been submitted to the PDB. All crystal structure figures in this paper were prepared using PyMol version 2.3 (Schrödinger LLC, Portland, OR).

**Table 1.** Crystallographic Data Collection and Refinement Statistics.

|  | CS | CS | PS F76A | PS F76A | PS F76A |
| --- | --- | --- | --- | --- | --- |
|  |  | 2FFPP |  | 2FFPP | DFFPP |
| *PDB ID* | xxxx | xxxx | xxxx | xxxx | xxxx |
| *Data collection statistics* |  |  |  |  |  |
| Space group | $P2_1$ | $P2_1$ | $P6_3$ | $P6_3$ | $P6_3$ |
| Wavelength (Å) | 0.9774 | 1.0 | 0.9792 | 0.9792 | 0.9792 |
| Resolution range (Å) | 90 - 2.33 | 20 - 2.65 | 20 - 2.50 | 20 - 2.20 | 20.0 - 2.65 |
| Highest resolution shell (Å) | 2.46 - 2.33 | 2.79 - 2.65 | 2.6 - 2.50 | 2.27 - 2.20 | 2.78 - 2.65 |
| Unit cell parameters (Å) | a = 68.5,<br>b = 89.8,<br>c = 114.1;<br>$\alpha = \gamma = 90$,<br>$\beta = 90.8$ | a = 68.7,<br>b = 89.6,<br>c = 114.8;<br>$\alpha = \gamma = 90$,<br>$\beta = 90.7$ | a = 182.4,<br>b = 182.4,<br>c = 56.3;<br>$\alpha = \beta = 90$,<br>$\gamma = 120$ | a = 181.5,<br>b = 181.5,<br>c = 55.8;<br>$\alpha = \beta = 90$,<br>$\gamma = 120$ | a = 180.5,<br>b = 180.5,<br>c = 55.9;<br>$\alpha = \beta = 90$,<br>$\gamma = 120$ |
| Total reflections | 388654 | 274946 | 770098 | 1119176 | 638633 |
| Unique reflections | 59306 | 40108 | 37349 | 53642 | 30519 |
| Completeness %<sup>a</sup> | 100 (100) | 99.0 (99.0) | 99.8 (100) | 99.9 (100) | 99.7 (100) |
| R<sub>merge</sub> %<sup>a</sup> | 20.4 (89) | 21.2 (100) | 9.8 (128) | 11.9 (186) | 11.1 (192) |
| CC(1/2)<sup>a</sup> | 1.0 (0.7) | 1.0 (0.7) | 1.0 (0.8) | 1.0 (0.8) | 1.0 (0.8) |
| I/σ (I)<sup>a</sup> | 7.5 (2.1) | 8.2 (2.2) | 20.0 (2.6) | 16.8 (2.1) | 18.9 (2.2) |
| Redundancy<sup>a</sup> | 6.6 (6.4) | 6.9 (6.9) | 20.6 (21.4) | 20.9 (21.3) | 20.9 (21.8) |
| *Refinement statistics* |  |  |  |  |  |
| Resolution range (Å) | 70 - 2.33 | 20 - 2.65 | 20 - 2.5 | 20.0 - 2.2 | 20 - 2.65 |
| No. of reflections used | 59266 | 40003 | 37331 | 53622 | 30479 |
| R<sub>cryst</sub> % | 21.0 | 20.4 | 21.1 | 18.5 | 19.8 |
| R<sub>free</sub> % | 25.6 | 24.9 | 23.9 | 21.3 | 22.9 |
| Protein atoms | 9381 | 9428 | 4732 | 5007 | 4927 |
| Ligand atoms | 52 | 139 | 20 | 26 | 26 |
| Metal atoms | 6 | 3 | 0 | 3 | 3 |
| Water molecules | 767 | 254 | 125 | 345 | 57 |
| r.m.s.d. in bond lengths (Å) | 0.002 | 0.001 | 0.001 | 0.002 | 0.002 |
| r.m.s.d. in bond angles (°) | 0.4 | 0.4 | 0.4 | 0.5 | 0.5 |

<sup>a</sup> Highest resolution shell values are given in parentheses.

## ACKNOWLEDGMENTS

We are grateful to the staff at the Advanced Light Source-Berkeley Center for Structural Biology for their assistance during X-ray data collection for CS datasets. The Advanced Light Source is funded by the Director, Office of Science, Office of Basic Energy Sciences, of the U.S. Department of Energy under Contract DE-AC02-05CH11231. The Berkeley Center for Structural Biology is supported in part by grants from the National Institute of General Medical Sciences.

The data set for PS F76A was collected at the Northeastern Collaborative Access Team beamlines, which are funded by the National Institute of General Medical Sciences from the National Institutes of Health (P30 GM124165). The Eiger 16M detector on the 24-ID-E beam line is funded by a NIH-ORIP HEI grant (S10OD021527). This research used resources of the Advanced Photon Source, a U.S. Department of Energy (DOE) Office of Science User Facility operated for the DOE Office of Science by Argonne National Laboratory under Contract No. DE-AC02-06CH11357.

This work was supported by National Institutes of Health grant T32GM007596 (J.O.M. and W.H.E.), RO1-GR403886 and RO1-GM096053 (D.L.T.), and the National Science Foundation CAREER program CHE-1253363 (I.J.K.).

## ABBREVIATIONS

FPP: farnesyl diphosphate

DFFPP: 12,13-difluorofarnesyl diphosphate

2FFPP: 2-fluorofarnesyl diphosphate

PS: pentalenene synthase

CS: caryolan-1-ol synthase.

## Notes

Funding: This work was supported by National Institutes of Health Grants T32GM007596 (J.O.M., W.H.E., M.P.) and the National Science Foundation CAREER program CHE-1253363 (I.J.K.).

### Competing Interest Statement

The authors have declared no competing interest.

### Summary of Updates

We wanted to include more citations.

